# Specific quorum sensing molecules are possibly associated with responses to herbicide toxicity in a *Pseudomonas* strain

**DOI:** 10.1101/2020.11.24.395699

**Authors:** Paloma Nathane Nunes de Freitas, Amanda Flávia da Silva Rovida, Caroline Rosa Silva, Sônia Alvim Veiga Pileggi, Luiz Ricardo Olchanheski, Marcos Pileggi

## Abstract

Pesticides contribute to pest control and increased agricultural production; however, they are toxic to non-target organisms and they contaminate the environment. The exposure of bacteria to these substances can lead to the need for physiological and structural changes for survival, which can be determined by genes whose expression is regulated by quorum sensing (QS). However, it is not yet clear whether these processes can be induced by herbicides. Thus, the aim of this work was to determine whether there is a QS response system in a *Pseudomonas fluorescens* strain that is modulated by herbicides. This strain was isolated from water storage tanks used for washing pesticide packaging and was tested against herbicides containing saflufenacil, glyphosate, sulfentrazone, 2,4-D, and dicamba as active molecules. We found that this strain possibly uses QS signaling molecules to control the production of reactive oxygen species, whether those produced by the bacterium’s energy generating system or by molecules induced by the presence of saflufenacil and glyphosate. This strain used other signaling molecules for various stages of biofilm formation in the presence of herbicides containing sulfentrazone, 2,4-D, and dicamba. These findings, as an initial screening which will guide new studies, suggest that this strain has a flexibility in gene expression that allows survival in the presence of several stress-inducing molecules, regardless of previous exposure. This represents a model of metabolic and physiological plasticity. Biofilms made up of several bacterial species can use this model in agricultural environments, increasing the potential for degradation of xenobiotics, but with impacts on diversity and functionality of microbiotas in these environments.

## 1. Introduction

Bacteria can change their behavior because of cell density, a phenomenon that mimics the behavior of multicellular organisms [1]. This is made possible through a bacterial communication system called quorum sensing (QS). This system allows bacteria to monitor the surrounding cell population density mediated by the production, release, accumulation and detection of chemical signaling molecules called autoinducers and, from there, they collectively coordinate gene expression, initiating important physiological changes for their survival [2], including bioluminescence, virulence, and biofilm formation [3]. In this way, bacteria can survive in environments where conditions constantly change, mediated by adaptations promoted by the flexibility of expression of their genes [4].

QS regulated the increase in the activities of antioxidative enzymes superoxide dismutase (SOD) and catalase (CAT) in *Salmonella typhi*, responding to reactive oxygen species (ROS) induced by bile, a bactericidal agent [5]. *Burkholderia cepacia,* an opportunistic bacterium associated with infections in individuals with cystic fibrosis, can evade the immune system and resist antibiotics with the help of virulence factors and biofilm formation, both of which are behaviors coordinated by QS [6]. Tang et al. [7] reported that the expression of CAT and the formation of biofilm, regulated by QS, contributed the ability of *Pseudomonas aeruginosa* PAO1 to resist nicotine stress. Via these mechanisms, several species of bacteria develop adaptive strategies coordinated by QS to enhance their survival in various stress conditions [8].

By contrast, little is known about the role of QS in bacterial survival in the context of environmental stresses. Bacteria need to deal with various toxic components in their environments, including pesticides used for pest control in the agriculture industry [9]. Although pesticide use is beneficial in this sense, there are also adverse effects of these molecules [10] in that they are also toxic to non-target organisms. Only 0.1% of pesticides reach target organisms, and the remainder contaminate the environment [11], being a major threat to biodiversity [12].

Thus, exposure of bacteria to toxic components can lead to changes in cell metabolism, growth, enzyme activity, and they can decrease the diversity of these organisms. Mazhari and Ferguson [13] found that glyphosate and paraquat herbicides led to decreased growth rates of bacterial strains isolated from herbicide degradation tanks. Meena et al. [14] reported that the herbicide 2,4-D inhibited the activities of *Rhizobium*, *Nitrosomonas*, and *Nitrobacter* spp., all of which are microorganisms important for the soil nitrogen cycle. Other studies have shown changes in the diversity of microbial communities after exposure to 2,4-D herbicides, atrazine, diuron [15], and mesotrione [16]. The latter induced oxidative stress in *Pantoea ananatis* CCT7673 [17] as well as changes in the pattern of lipid saturation in *Bacillus megaterium*, with possible changes in membrane permeability [18].

It remains unclear as to whether QS can be used by bacteria to coordinate population behavior in response to herbicides. Therefore, the objective of this work was to determine whether there is a specific QS-mediated herbicide response to ROS induced by herbicides in a *Pseudomonas fluorescens* strain isolated from an environment contaminated with pesticides.

## 2. Material and methods

The experimental design (Fig. 1) included a bacterial strain and five different herbicides. The strain was subjected to combinations of treatments and bacterial growth conditions to investigate a QS response system model modulated by the presence of herbicides. Five aspects were investigated: bacterial growth curve, cell viability, extraction of QS signaling molecules [analyzed using liquid chromatography coupled to mass spectrometry (LC-MS/MS)], biofilm formation analysis, and statistical analysis for each stage.

**Figure 1.**
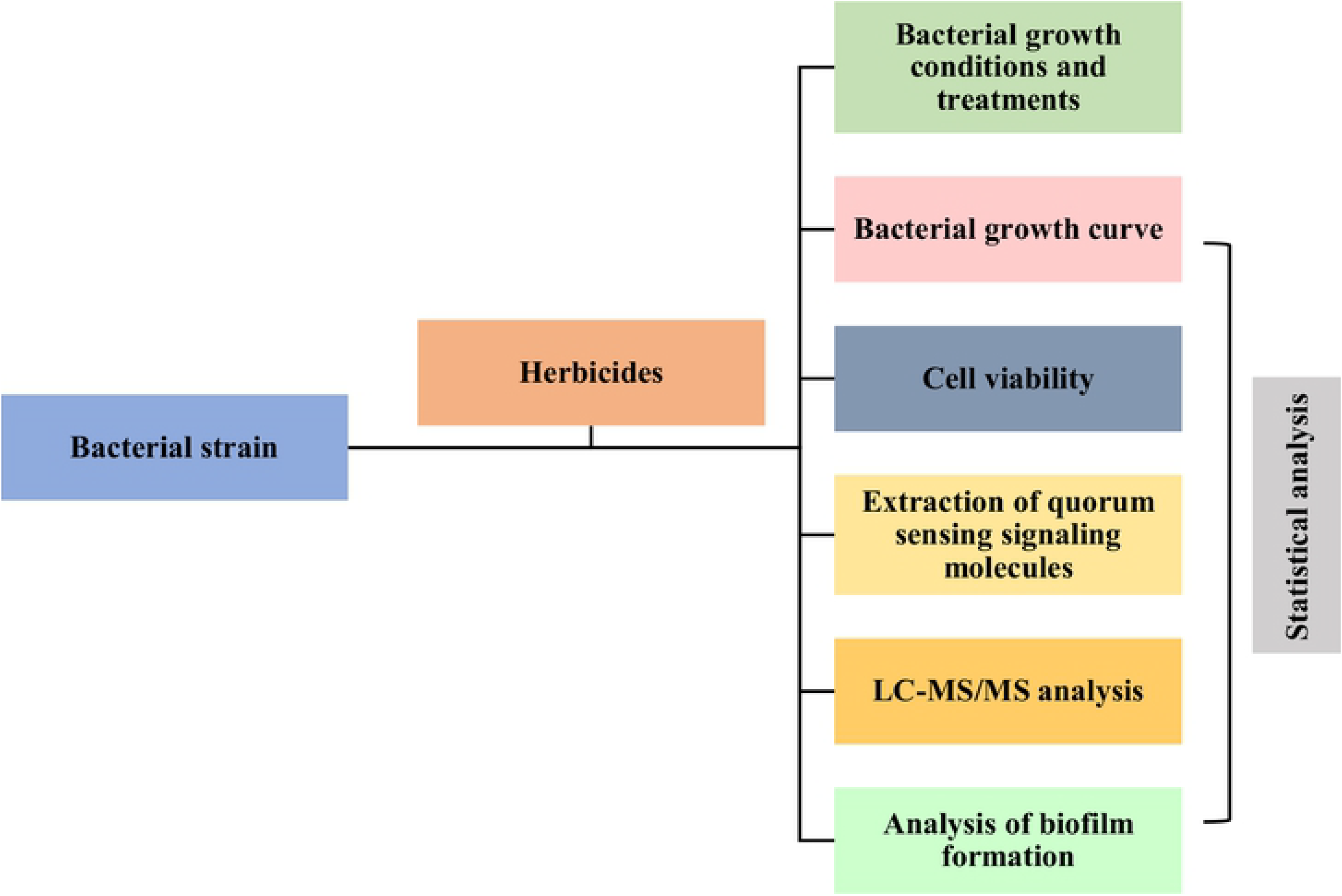
Experimental design of the study.

Experimental design

### 2.1 Bacterial strain

The strain studied here was obtained from water storage tanks used to wash pesticide packaging, belonging to BASF Research Station, located at Fazenda Escola Capão da Onça of the State University of Ponta Grossa (UEPG). The list of pesticides, whose packaging has been washed, was provided by BASF (Ludwigshafen on the Rhine, Rhineland-Palatinate, Germany) [19]. The strain was identified by sequencing the amplicon 16S RNA, displaying 98.35% identity with *Pseudomonas fluorescens*, accession number **KY807296** - NCBI, identified as *P. fluorescens* CMA55. The strain was deposited at the Johanna Döbereiner Biological Resources Center (CRB-JD) - Embrapa Agrobiologia, under the code **BR14566** and is part of the Collection of Environmental Microorganisms of the UEPG’s Laboratory of Environmental Microbiology.

### 2.2 Herbicides

The herbicides studied in this work are shown in Table 1.

**Table 1.** Trademark, registration company and active ingredient of the herbicides used.

| Trademark | Registration company | Active ingredient |
| --- | --- | --- |
| Heat | BASF | Saflufenacil (N'-{2-chloro-4-fluor-5-[1,2,3,6-tetrahydro-3-methyl-2,6-dioxo-4-(trifluoromethyl)pyrimidine-1-yl]benzoyl}-N-isopropyl-N-methylsulfamide) |
| Boral 500 SC | FMC (Philadelphia, PA, EUA) | Sulfentrazone (2',4'-dichloro-5'-(4-difluoromethyl-4,5-dihydro-3-methyl-5-oxo-1H-1,2,4-triazol-1-yl) (methanesulfonanilide) |
| Roundup Transorb R | MONSANTO (Saint Louis, MO, EUA) | Glyphosate (potassium salt of (N-(phosphonomethyl)glycine) |
| Aminol 806 | ADAMA (Raleigh, NC, EUA) | 2,4-D (dimethylamonia (2,4-dichlorofenoxy) acetate (2,4-D dimethylamine) |
| Atectra | BASF | Dicamba (3,6-dichloro-o-anisic acid) |

Herbicides information

### 2.3 Bacterial growth conditions and treatments

For the growth of *P. fluorescens* CMA55, we used culture conditions designed for the growth of this strain, with mineral medium (MM - 35,2 g L^−1^ KH_2_PO_4_; 53 g L^−1^ Na_2_HPO_4_; 1,696 g L^−1^ Mn SO_4_; 2,7136 g L^−1^ CaCl_2_; 0,0672 g L^−1^ FeSO_4_; 0,0104 g L^−1^ Na_2_MoO_4_; 0,08 g L^−1^ CuSO_4_; 1,24 g L^−1^ H_3_BO_4_; 0,25 g L^−1^ ZnSO_4_; 0,0056 g L^−1^ CoCl_2_; 2,5 g L^−1^ (NH_4_)_2_SO_4_; 3 g L^−1^ MgSO_4_), adding 50 g L^−1^ sucrose and 50 g L^−1^ glucose, according to Gädke et al. [20]. For the treatments, MM was supplemented with herbicides, all water soluble, as follows: Heat (H – 0.49 mM), Boral 500 SC (B – 0.049 mM), Roundup Transorb R (R – 0.233 mM), Aminol 806 (A – 0.173 mM), and Atectra (At – 0.103 mM). The concentrations of herbicides are equivalent to the dose used in agriculture. The growth conditions were 30 °C and under agitation at 120 rpm. The experiments were carried out in triplicate.

### 2.4 Bacterial growth curve

The bacterial strain was grown for 24 h in MM (pre-inoculum). The inoculum was assayed in the treatments from the pre-inoculum at an initial optical density (OD) of 0.05 to 600 nm. Bacterial growth was monitored every 1 h in a spectrophotometer at 600 nm.

### 2.5 Cell viability

The same pre-inocula and conditions were used as previously described (sections 2.3 and 2.4), with 100 μL aliquots being removed after 14 h of incubation (established as the final period of the log phase and the beginning of the stationary growth phase). The samples were diluted to 10^−7^, seeded in MM with agar (20 g L^−1^) and incubated at 30 °C. After 24 h, the colony forming units (CFU) were counted.

### 2.6 Extraction of quorum sensing signaling molecules

The extraction of QS molecules was performed using the liquid-liquid method [21] at 14 h of bacterial growth. In addition to the treatments described in the section 2.3, medium 0 (MM without herbicide and without bacteria) was used as control. A total of 100 ml of culture medium were removed, centrifuged (8.000 x *g*) for 5 min, 2.0 ml of the supernatants were collected, and 10 μM of the internal standard N-heptanoyl-L-homoserine lactone (C7-HSL) was added. The extraction process was carried out using ethyl acetate, according to Ortori et al. [21] with modifications. A total of 2.0 ml of ethyl acetate acidified with 0.01% acetic acid were added. The solution was stirred for 1 min, and the organic phase was collected. The extraction procedure was repeated twice, and the extracts were evaporated in an exhaust hood at room temperature. The dry samples were stored at –80 °C until use, at which time they were reconstituted in 100 μL of methanol (MeOH) immediately before analysis using LC-MS/MS.

### 2.7 LC-MS/MS analysis

The identification analyses of N-acyl homoserines lactones (AHL) and 2-alkyl-4-(1H)-quinolones (AQ) were carried out according to the method described by Ortori et al. [21], with modifications. A total of 10 μL of the extracts prepared at a flow rate of 0.3 mL/min and were injected into the LC-MS/MS model Acquity and Xevo TQD equipment (Waters Corporation, Milford, MA, USA) equipped with binary pumps, a vacuum degasser, a SIL-HTc automatic sampler and quadrupole 4000 triple linear ion traction mass spectrometer equipped with a Turbo-Ion source. The column used was an ACQUITY UPLC® BEH C18 2.1 × 50 mm (1.7-μm porosity), maintained at 50 °C. The UPLC system consisted of 0.1% formic acid (mobile phase A) in ultrapure water and a mobile phase B consisting of 0.1% formic acid (Fisher Scientific, Loughborough, UK) in acetonitrile. The gradient profile was as follows: isocratic over 1 min, a linear gradient of 50% A and 50% B over 1.5 min, then an additional gradient of 1% A and 99% B over 5.5 min followed by 1% A and 99% B for 7 min and the condition was restored to 90% A and 10% B for 10 min. The column was rebalanced for a total of 2 min. Mass spectrometry (MS) experiments were conducted in the positive ionization mode in electrospray ionization (ESI). The voltage of the capillary cone was 4 kV, the temperature of desolvation was 450 °C and the volume of desolvation was 550 L/Hr. S1 Appendix describes the analytical parameters unique to each analyte, including cone voltage and collision energy, in addition to the precursor and child ions used to configure the multiple reaction monitoring (MRM) mode for each individual analyte. The optimization of the analytical parameters in MS was performed by infusing the internal standard N-heptanoyl-L-homoserine lactone (C7-HSL) (Sigma-Aldrich, Saint Louis, MO, USA).

### 2.8 Analysis of biofilm formation

Using the same conditions of pre-inoculum and inoculum described in the section 2.4, the formation of biofilm in microplate was evaluated using the crystal violet (CV, 0.1%) staining method, according to Tang et al. [7], with modifications. After 14 h of incubation, the culture medium was removed and the cells adhering to the surface were stained for 30 min with CV. The wells were washed three times with distilled water to remove the unabsorbed CV, and then the adherent CV was solubilized with 95% alcohol and measured at 570 nm using a microplate reader (Elx808™, BioTek, Winooski, VT, USA).

### 2.9 Statistical analysis

Statistical analyses were performed using the Tukey test and the significance criterion p <0.05 using software R version 3.5.1. The identified molecules were analyzed using PCA, a multivariate analysis technique that reduces many original correlated variables to a small number of new and unrelated variables [22].

## 3. Results and discussion

### 3.1 QS signaling molecules and population density in *Pseudomonas fluorescens* CMA55

The bacterial communication system depends on a population cell density to produce and identify signals at levels sufficient for gene regulation. Many QS-dependent genes are activated after the middle or late exponential growth phase and not during earlier growth phases [23]. Others are activated during the stationary growth phase [24]. Therefore, as an initial screening searching for herbicides responses, we chose 14 h for the extraction and analysis of QS signaling molecules. This point was identified from the growth curve of *P. fluorescens* CMA55 (Fig. 2) as the final period of the log phase and beginning of the stationary growth phase.

**Figure 2.**
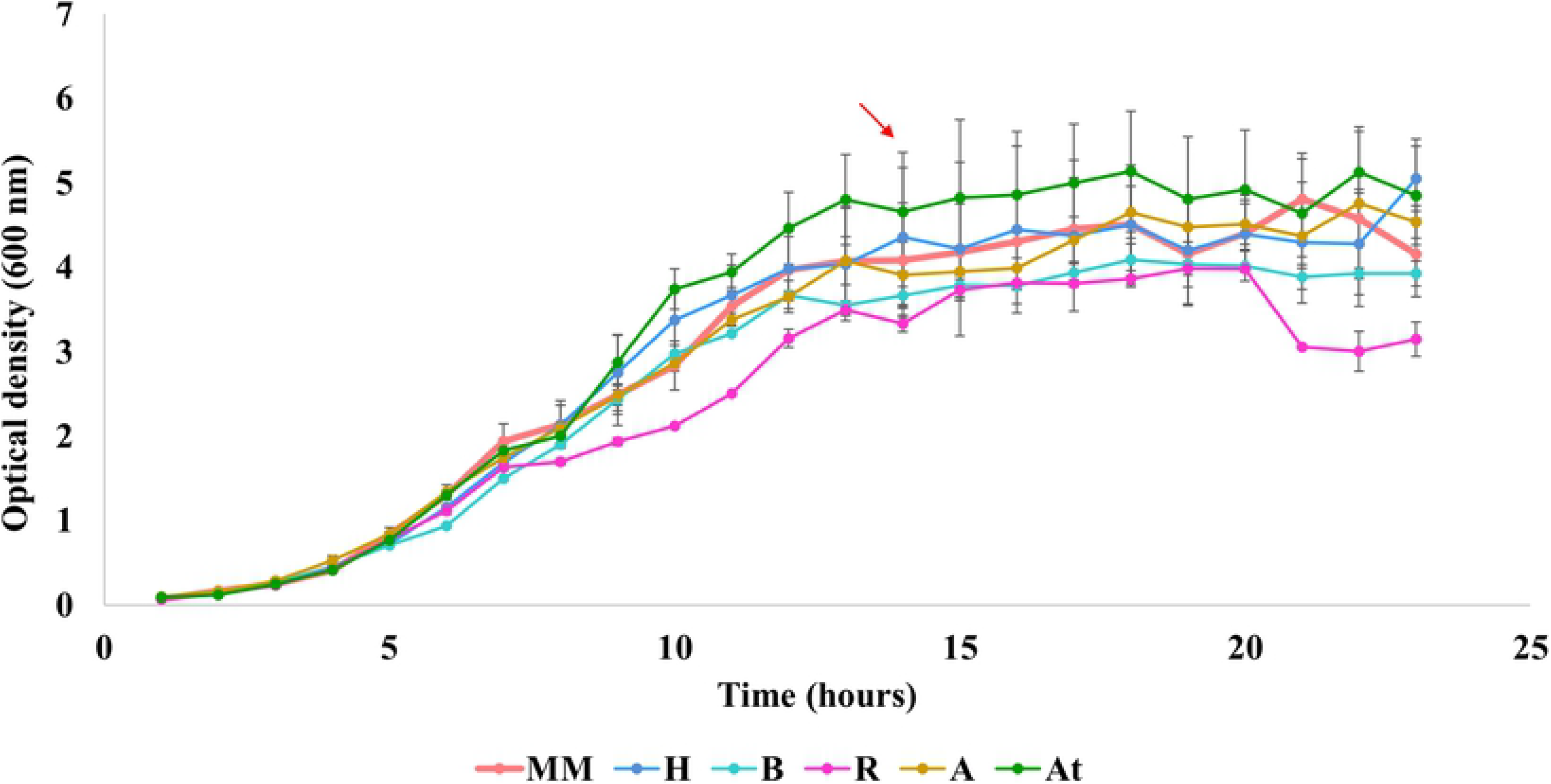
Growth curves of the *Pseudomonas fluorescens* CMA55 strain in control (MM) and in the Heat (H), Boral 500 SC (B), Roundup Transorb R (R), Aminol 806 (A) and Atectra (At) treatments. The strain reaches the final period of the log phase and the beginning of the stationary growth phase at the 14 h time period (arrow). The bars represent the standard errors in the averages. (p <0.05).

Growth curves

### 3.2 Viability of *Pseudomonas fluorescens* CMA55 against herbicides

The tolerance data for Heat, Boral 500 SC, Roundup Transorb R, Aminol 806, and Atectra herbicides presented by the microorganisms isolated from water storage tanks used for washing pesticide packaging, including *P. fluorescens* CMA55, are in Fig. 3. These data show that the number of tolerant organisms decreases as the concentration of herbicides increases, primarily with respect to Roundup Transorb R. Dennis et al. [25] reported that glyphosate, glufosinate, paraquat, and paraquat-diquat presented little threat to biodiversity or the function of the soil microbial community. However, the same authors suggested that the organisms that were not tolerant to herbicides had already disappeared; therefore, it only appeared that there was no impact of herbicides on the soil community. We believe that the disappearance of non-tolerant species is a threat to biodiversity. Exposure of toxic substances imposes selective pressure on sensitive species that subsequently disappear, leaving tolerant species to dominate the community [26].

**Figure 3.**
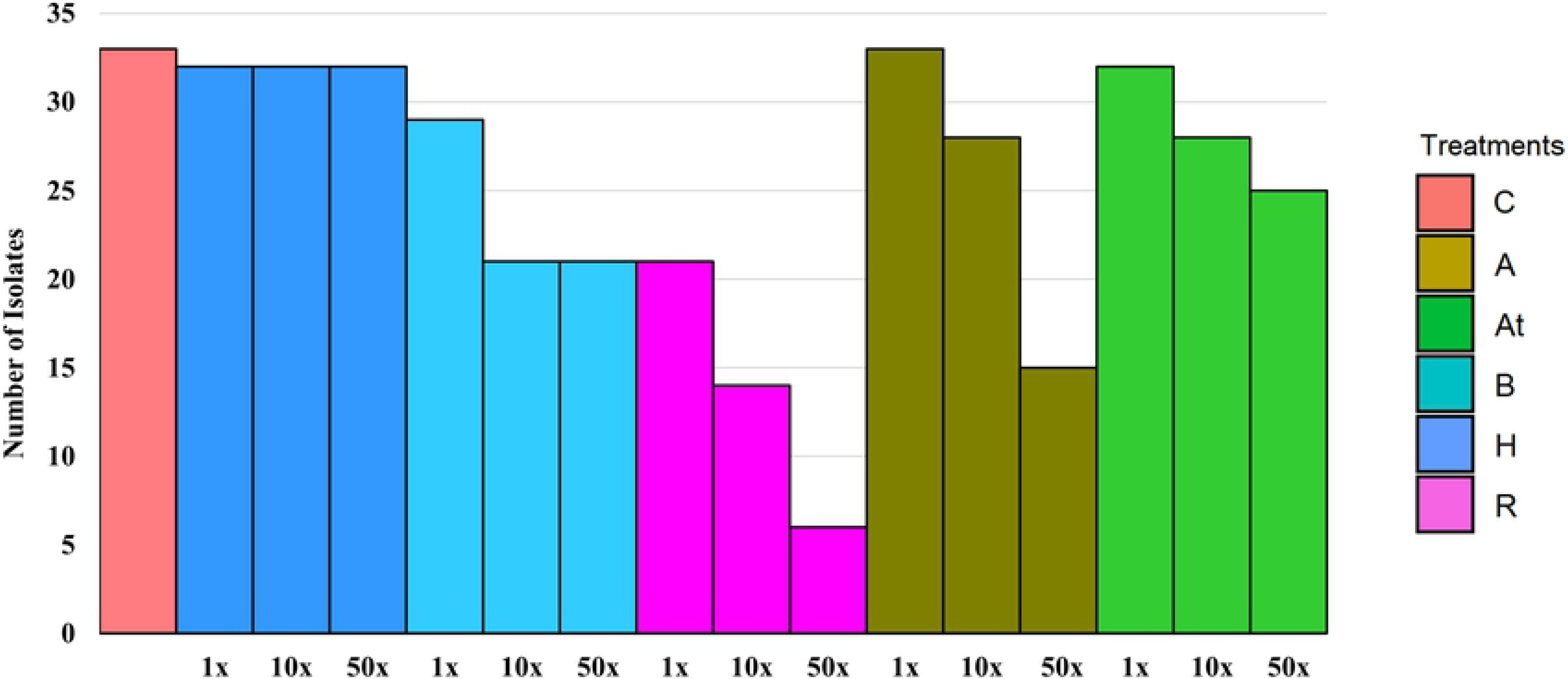
Isolates from water storage tanks used for washing pesticide packaging that showed tolerance in the three concentrations (1x, 10x and 50x the field dose used in agriculture) in the treatments Heat (H), Boral 500 SC (B), Roundup Transorb R (R), Aminol 806 (A) and Atectra (At).

Tolerance to herbicides by isolates from the water collection

Other studies show that herbicides can modify the structure of the microbiota; these include chlorimuron-ethyl and metsulfuron-methyl herbicides that induced a bacteriostatic effect in *Burkholderia pseudomallei*, *P. aeruginosa*, and *Acinetobacter baumannii* [27]. These authors also showed that *Pseudomonas putida*, *Escherichia coli*, and *Salmonella typhimurium* were inhibited at low concentrations of glyphosate, because they possessed 5-enolpyruvylshikimate-3-phosphate synthase (EPSPS). The glyphosate herbicide reduced the growth and performance of *Rhizobium*, *Burkholderia*, and *Pseudomonas* spp., as well as those of arbuscular mycorrhizal fungi [28]. This 2,4-D herbicide altered the surface, roughness, and surface load of the bacterial cell envelope of *E. coli*, leading to loss of structure integrity, allowing entry of the herbicide and consequently leading to the production of ROS [29]. The functionality of mycorrhizal fungi and rhizobia in soy was reduced using the sulfentrazone herbicide [30]. De Novais et al. [31] showed a significant negative impact of the herbicide dicamba on arbuscular mycorrhizal fungi. There are few published data on the toxicity of the herbicide saflufenacil in bacteria.

The data of the growth curve (Fig. 2) and cell viability (Fig. 4) of *P. fluorescens* CMA55 showed no significant differences in the presence of the tested herbicides with respect to control at 14 h, suggesting tolerance. Probably at this time, a population density was reached in which changes in gene expression, and consequently in the physiology of cells, were triggered [32], contributing to herbicide tolerance.

**Figure 4.**
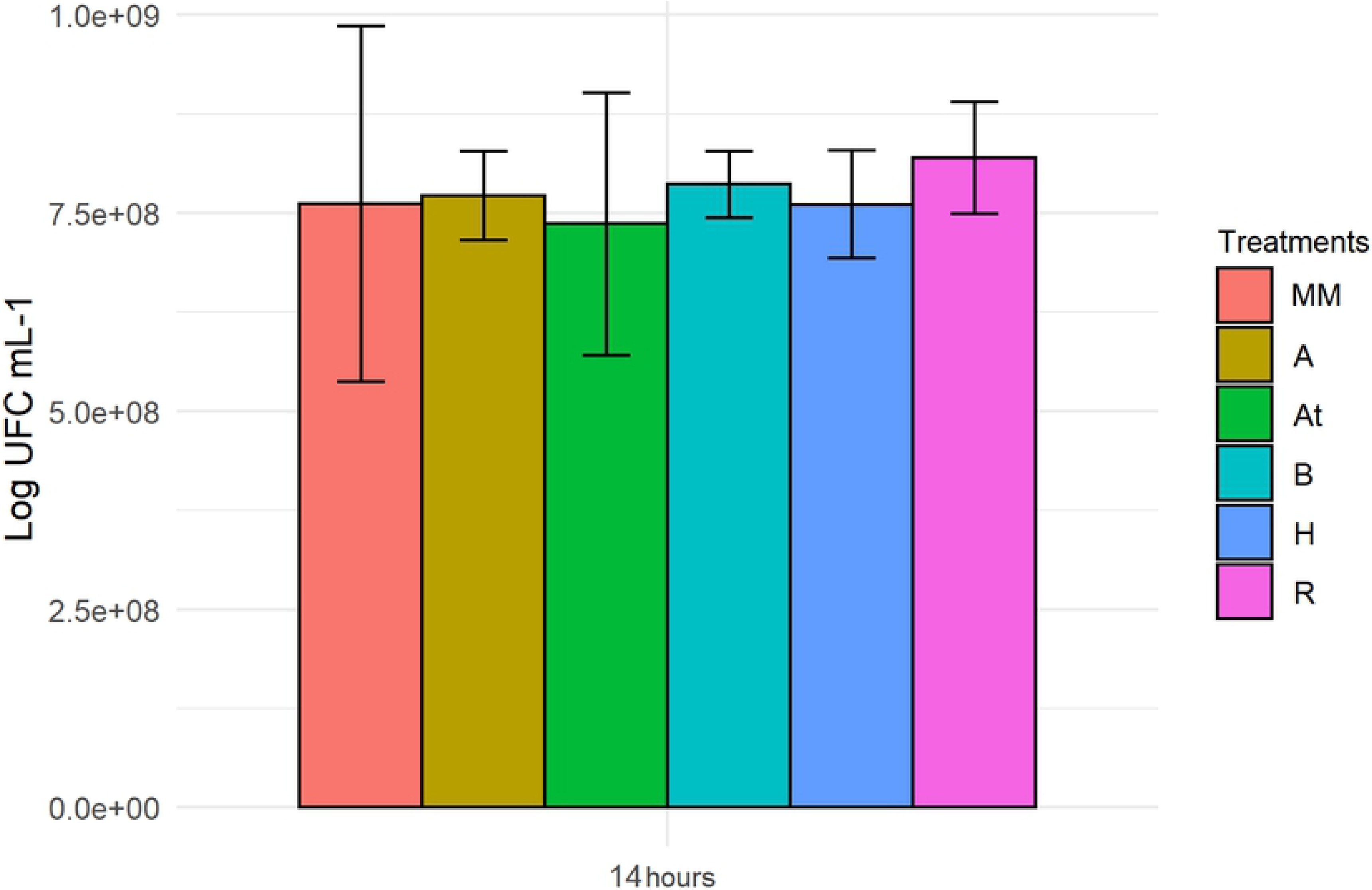
Cell viability of the *Pseudomonas fluorescens* CMA55 strain after 14 h of incubation in control (MM) and in the Heat (H), Boral 500 SC (B), Roundup Transorb R (R), Aminol 806 (A), and Atectra (At). During this period of growth, there were no significant differences in treatments compared to control. The bars represent the standard errors in the averages. (p <0.05).

Cell viability

### 3.3 Profile of QS signaling molecules in response to herbicides

The linking of specific treatments with 22 QS signaling molecules identified in this work was performed using PCA. In controls 0 and MM and in treatments H (Heat) and R (Roundup Transorb R), there were similar profiles of QS molecules (Fig. 5). However, in treatments B (Boral 500 SC), A (Aminol 806), and At (Atectra), there was another profile of molecules that was similar between these treatments but distinct from controls 0 and MM.

**Figure 5.**
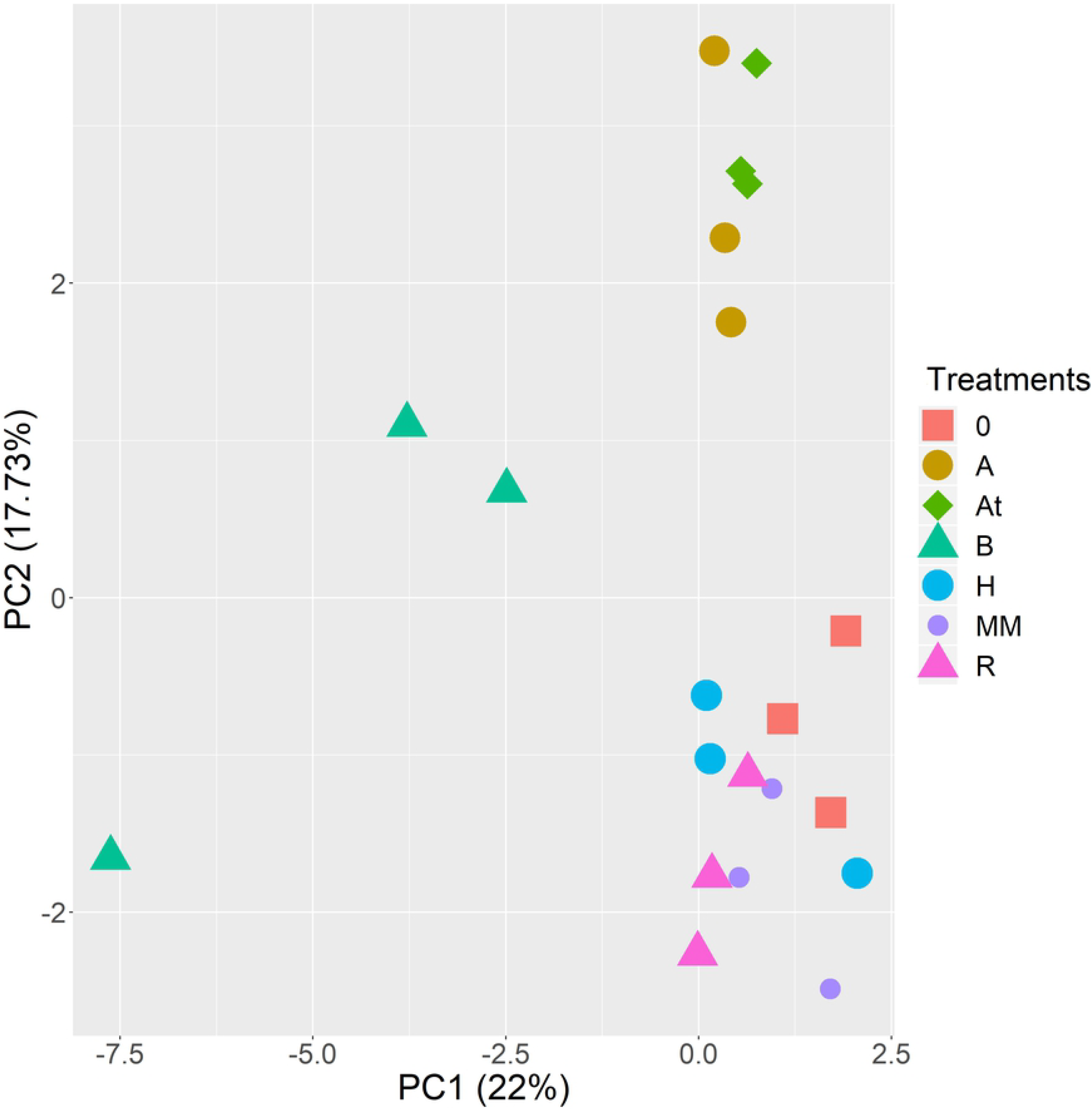
PCA analysis on the profile of molecules signaled by *Pseudomonas fluorescens* CMA55. The treatments (in triplicates) are represented by the symbols (controls 0 and MM and treatments H - Heat, B - Boral 500 SC, R - Roundup Transorb R, A - Aminol 806 and At - Atectra). The explanation percentages are PC1 = 22 and PC2 = 17.73.

PCA analysis

Most of the QS molecules in controls 0 and MM and in the H (Heat) and R (Roundup Transorb R) treatments (Table 2) are related to stress, including C8-HSL, 3-oxo-C8-HSL and 3-OH-C8-HSL. In *B. pseudomallei*, these molecules are related to the expression of the *dpsA* gene, a regulator of catalase and peroxidase activities that play fundamental roles in protecting bacterial DNA from oxidative stress [33]. In *Salmonella enteritidis* PT4 578, the presence of the C12-HSL molecule increased the abundance of thiol proteins, suggesting that cells can prepare for possible oxidative stress, demonstrated by the prevention of structural changes caused by ROS by the SdiA protein [34]. The molecules 3-oxo-C6-HSL and HQNO are involved in virulence factor behaviors [35] and the 3-OH-C4-HSL molecule appears to be involved in various stages of biofilm formation [36]. No data were found in the literature regarding the function of this group of molecules in the genus *Pseudomonas*.

**Table 2.** Area and retention time of the QS signaling molecules in controls 0 and MM and in treatments H and R, as well as some functions of these molecules that have already been described in the literature.

| Molecule | Area | Retention time | Functions |
| --- | --- | --- | --- |
| C8-HSL | 756.88 | 6.75 | Oxidative stress |
| C12-HSL | 9190.68 | 6.36 | Oxidative stress |
| 3-oxo-C6-HSL | 306.425 | 0.76 | Virulence factors |
| 3-oxo-C8-HSL | 563.005 | 7.37 | Oxidative stress |
| 3-OH-C4-HSL | 1252.065 | 4.39 | Biofilm |
| 3-OH-C8-HSL | 135.79 | 0.56 | Oxidative stress |
| HQNO | 35.21 | 7.15 | Virulence factors |

Functions of QS related to controls, H and R herbicides treatment

Because most of these molecules are related to stress, our hypothesis was that the antioxidant system of *P. fluorescens* CMA55 is under regulatory control of QS. Bollinger et al. [37], reported that responses to oxidative stress are regulated by QS, including the enzymes Mn-SOD and KatA. The similarity of molecules in treatments and controls may be due to the use of this stress regulation system, because ROS participate in normal aerobic metabolism; however, their levels can be increased by exogenous sources such as heavy metals, UV radiation, and environmental pollutants [38]. In this way, the strain may exploit the same defense mechanism against ROS from different sources.

Regarding the profile of QS molecules in treatments B (Boral 500 SC), A (Aminol 806), and At (Atectra), these molecules were related to adaptive and bacterial survival strategies via virulence factors and biofilm formation (Table 3).

**Table 3.** Area and retention time of the QS signaling molecules in treatments B, A, and At, as well as some functions of these molecules that have already been described in the literature.

Functions of QS related to B, A and At herbicides treatments
| Molecule | Area | Retention time | Functions |
| --- | --- | --- | --- |
|  |  |  | Biofilm Formation / Virulence Factor* |
| C4-HSL | 48738.8 | 6.35 | Maturation |
| C6-HSL | 279.76 | 3.23 | Maturation |
| C10-HSL | 180.035 | 4.96 | Initial adhesion |
| C14-HSL | 6532.94 | 7.03 | SPE matrix |
| 3-oxo-C4-HSL | 5871.44 | 5.16 | Unknown |
| 3-oxo-C10-HSL | 528.195 | 2.78 | Unknown |
| 3-oxo-C12-HSL | 1293.435 | 6.50 | Initial adhesion |
| 3-oxo-C14-HSL | 600.655 | 7.36 | SPE matrix |
| 3-OH-C10-HSL | 426.895 | 3.77 | SPE matrix |
| 3-OH-C12-HSL | 35560.6 | 4.56 | SPE matrix |
| PQS | 208.265 | 6.16 | SPE matrix and Maturation<br>Production of pyocyanin, ramnolipids and regulation of efflux pumps* |
| C9-PQS | 2093.51 | 3.48 | Maturation<br>Production of pyocyanin, ramnolipids and regulation of efflux pumps* |
| C11-PQS | 225.585 | 3.13 | Maturation<br>Production of pyocyanin, ramnolipids and regulation of efflux pumps* |
| UHQ | 62.08 | 3.45 | Maturation<br>Production of pyocyanin, ramnolipids and regulation of efflux pumps* |
| NQNO | 21778.5 | 3.48 | Maturation<br>Production of pyocyanin, ramnolipids and regulation of efflux pumps* |

Note: (\*) Molecule functions PQS, C9-PQS, C11-PQS, UHQ and NQNO – Virulence Factor.

Functions of QS related to B, A and At herbicides treatments

Wang et al. [39] observed that C10-HSL is one of the main AHL involved in the biofilm development process that is related to the initial fixation process of this structure in response to environmental factors (Table 3). The molecule 3-oxo-C12-HSL is also associated with the initial phase of biofilm development in *P. aeruginosa* PAO1, which is fundamental for the pathogenicity of this bacterium [40]. Mukherjee and Bassler [2] reported that *P. aeruginosa* uses QS to coordinate the formation of biofilms, contributing to its ability to persist and progress because this structure allows resistance to the action of antibiotics in human infections. In this manner, the signaling mediated by 3-oxo-C12-HSL and C10-HSL suggests that *P. fluorescens* CMA55 also initiates biofilm formation to increase the chance of survival in the presence to stress induced by the herbicides Boral 500 SC, Aminol 806, and Atectra.

PQS is involved in the formation of one of the components of the matrix of extracellular polymeric substances (SPE), extracellular DNA (eDNA), which acts as a structural support to maintain the biofilm architecture in *P. aeruginosa* [41]. In addition, PQS and molecules such as C9-PQS, C11-PQS, UHQ, and NQNO are involved in the production of pyocyanine [42]. eDNA and pyocyanine are essential for biofilm maturation, because they bind and increase biofilm viscosity, influencing the physical-chemical interactions of the biofilm matrix with the environment, thereby facilitating cell aggregation [43]. The SPE matrix acts as a physical barrier, preventing toxic agents from reaching their sites of action, including the outer membrane in gram-negative bacteria [44]. In addition, the matrix interacts and absorbs herbicides [26]. Thus, the relationship of the PQS, C9-PQS, C11-PQS, UHQ and NQNO molecules in the presence of Boral 500 SC, Aminol 806, and Atectra may indicate that *P. fluorescens* CMA55 is using the biofilm to provide protection against the toxicity of these herbicides.

The PQS, C9-PQS, C11-PQS, UHQ, and NQNO molecules are also involved in the production of ramnolipids, which contribute to the formation of internal cavities within the mature biofilm, allowing the flow of water and nutrients [45] and in the regulation of efflux pumps, multiprotein complexes that cross the envelope of gram-negative bacteria, responsible for expelling various toxic components and a wide variety of antimicrobials [46]. The relationship of efflux pumps with herbicide tolerance, as described in Kurenbach et al. [47], may be occurring in *P. fluorescens* CMA55 in response to Boral 500 SC, Aminol 806, and Atectra. Regulation of this system by QS as a bacterial response mechanism to environmental stresses is a possibility that should be further investigated.

Biofilm formation in a *Salmonella* strain is influenced by nutrient deprivation conditions, when it produces less SPE than under cultivation conditions in more nutrient-rich culture media [48]. The molecules C14-HSL, 3-oxo-C14-HSL, 3-OH-C10-HSL, and 3-OH-C12-HSL are associated with the biofilm matrix of *Acidithiobacillus ferrooxidans*. In this bacterium, the quantity of lipopolysaccharides that constitute the matrix increases with the decrease of the phosphate content in culture media [49]. The signaling from these same molecules in *P. fluorescens* CMA55 suggest that the SPE matrix mediates metabolic flexibility, permitting survival in environments with unfavorable nutritional conditions, including mineral environments, in addition to the presence of herbicides. These data suggest that nutritional stress may be one of the primary mechanisms responsible for the aggregation and formation of biofilms, representing an adaptive to unfavorable environments.

The molecules C4-HSL and C6-HSL are important for biofilm maturation processes in *P. aeruginosa* PAO1, for the formation of the SPE matrix, and for channels for transporting water and nutrients [40]. These channels are like a circulatory system, distributing various nutrients and removing biofilm residues [50]. As the biofilm matures, bacteria become more resistant to environmental stresses or antibiotics [51]. The same role could be played by these molecules in *P. fluorescens* CMA55 to increase resistance to herbicides.

There are few references in the literature regarding 3-oxo-C4-HSL and 3-oxo-C10 HSL (Table 3). Hansen et al. [52] showed that, in *Aeromonas salmonicida*, there is production of these molecules; however, it remains unknown what behaviors are activated by them. They are probably associated with virulence behavior or biofilm formation.

The QS system mediated by AHL and AQ in gram-negative bacteria such as *Pseudomonas* is involved in behaviors such as virulence and biofilm formation, and may confer resistance against antimicrobials, phagocytosis, oxidative stress, and lack of nutrients, among others, allowing survival in various environmental niches [8]. Possibly, the 15 molecules described indicate similar roles *P. fluorescens* CMA55 in the presence of the herbicides Boral 500 SC, Aminol 806, and Atectra. However, this work is an initial screening which will guide new studies of phenotype complementation (tolerance to herbicides, response to oxidative stress and biofilm production) with mutants lacking the production of QS molecules evaluated in this work, searching for the transcriptional regulator that interact with each of these quorum-sensing related molecules and the target genes that are regulated by the transcriptional regulator-QS complex.

### 3.4 Biofilm formation by *Pseudomonas fluorescens* CMA55

The formation of biofilm is a gradual process in which planktonic bacterial cells, upon reaching a certain density, acquire the potential to live as multicellular entities [53]. It has been reported that this process is regulated by the QS [54]. The process involves the secretion of a matrix composed of SPE responsible for the adhesion of planktonic bacteria on a surface, followed by the formation of microcolonies, then the development of the architecture and maturation of the biofilm. When the environment is no longer favorable to its maintenance, there is the dispersion of bacteria that can colonize new environments, restarting the process [50].

In our study, three significantly different groups were found, in decreasing order of biofilm production in the 14 h period (Fig. 6). The first is composed of control, in which MM, despite presenting consistent growth rates (Fig. 2), may have caused some type of nutritional stress as it is a medium with defined composition, and therefore induced the formation of biofilm [55]. The second group contains treatments B, A, and At, for which it was already demonstrated that QS molecules signal biofilm formation (Table 3), a hypothesis confirmed with the results shown in Fig. 6. These results suggest that biofilms increase the adaptive repertoire of the strain in the context of unfavorable nutritional conditions, in the presence of herbicides in vitro, and possibly in isolation environments. Higher biofilm production in nutrient-deprived culture media in bacteria such as *Salmonella* suggest the influence of nutritional conditions on regulation in these structures [48]. Biofilm formation can be considered a bacterial adaptive strategy in that it permits survival compared with planktonic forms in the form of resistance to environmental stresses [56]. In addition, biofilm can be a sustainable alternative in the recovery of environments contaminated by toxic pollutants [54].

**Figure 6.**
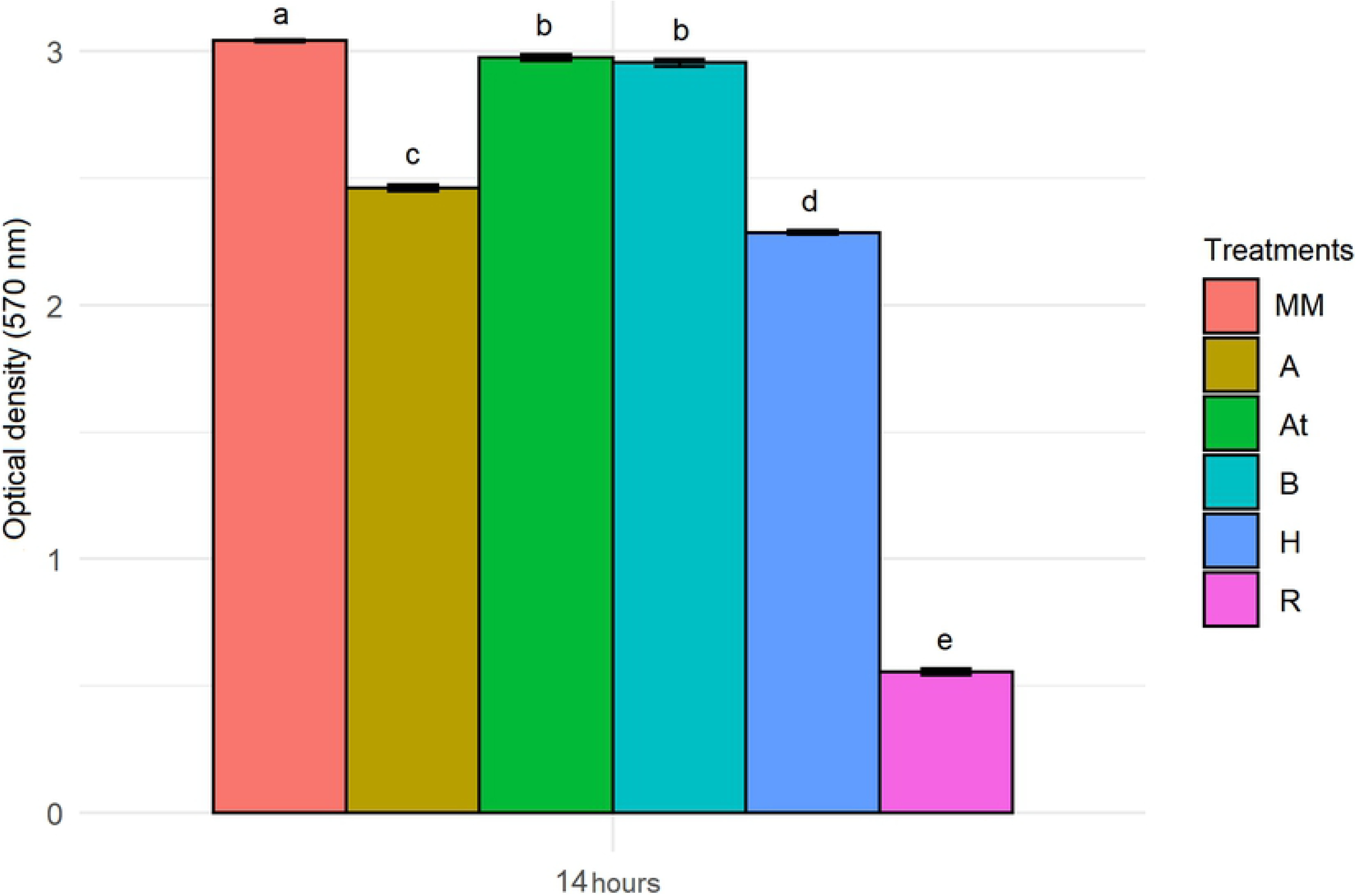
Quantification of biofilm from the *Pseudomonas fluorescens* CMA55 strain after 14 h of incubation in control (MM) and in the Heat (H), Boral 500 SC (B), Roundup Transorb R (R), Aminol 806 (A), and Atectra treatments (At). The bars represent the standard errors in the averages. The letters (a, b, c, d, e) represent a degree of significance between treatments. (p <0.05).

Quantification of biofilm

The third group consists of treatments H and R, in which only one of the QS molecules, 3-OH-C4-HSL, which has also been identified in MM, signals biofilm formation. The other molecules identified in these treatments with these herbicides signal the activation of processes that combat stress (Table 2), indicating a change in response strategy for treatments H and R in relation to B, A, At, and MM.

Metabolic and physiological plasticity allows bacteria to engage different adaptation response mechanisms depending on environmental conditions. In *P. aeruginosa*, a versatile metabolism associated with QS appears to be essential for processes such as invasion of new niches, acquisition of nutrients, microbial competition, and virulence [57]. Sudden variations in temperature did not affect the survival capacity of several bacterial genera, including *Pseudomonas*, mediated by an increase in unsaturated fatty acids and the production of QS-regulated zeaxanthin pigments, all of which are vital characteristics that stabilize the fluidity of membranes and mediate cryoprotection. These physiological changes were important for their survival, and could be explained by physiological plasticity [58].

It may be that the *P. fluorescens* CMA55 strain used biofilm production strategies and activation of antioxidative enzymes under the direction of QS molecules in response to both herbicides for which there was previous contact (Heat, Aminol 806 and Atectra) and for those that did not exist (Roundup Transorb R and Boral 500 SC), in the water storage tanks used to wash pesticide packaging. This would suggest, therefore, that this strain shows flexibility in gene expression associated with QS that permits survival in the context of stress-inducing molecules present in herbicides, regardless of previous contact, and may represent a model of metabolic and physiological plasticity necessary for adaptation to agricultural environments. These issues can be better elucidated with mutational complementation studies, which can be drawn from this work. Due the glyphosate toxicity to bacterial strains (Fig. 3) the study of the inhibition of quorum-sensing system, or quorum quenching, involved in biofilm formation by this herbicide is the next logical step.

The QS regulates important bacterial behaviors and is considered a very promising field of study for several areas. Pieces of information about chemical and biological interactions of bacterial communities in environments in constant exposure to pesticides can be analyzed through bioinformatics and analytical techniques. It is essential to consider that little is known about the ecological role of QS in the adaptation and survival of bacteria under environmental stresses, especially herbicides.

## 4. Conclusions

We described the relationship between QS signaling molecules and herbicides in a bacterial strain, *P. fluorescens* CMA55, isolated from water storage tanks used to wash pesticide packaging. No reports have yet been found in the literature on this approach. This strain possibly uses QS signaling molecules to control the production of ROS. Regardless of whether they are produced by the bacterium’s energy generating system or whether they are induced by the presence of the herbicides Heat and Roundup Transorb R, the latter was not present at the isolation site of the bacterium, suggesting that the organism developed adaptive potential for stressful conditions of a different sort. On the other hand, the same bacterium presented a modulated response system against the herbicides Boral 500 SC (not present in the isolation site), Aminol 806, and Atectra, using other signaling molecules involved in different stages of biofilm formation, suggesting flexibility in gene expression that allows survival in the context of different stress-inducing molecules, but using universal strategies such as those evolved for structuring biofilms.

The *P. fluorescens* CMA55 strain used biofilm production strategies and activation of antioxidative enzymes under signaling of QS molecules to respond to herbicides, irrespective of previous contact. These findings suggest that the organism has flexibility with respect to gene expression associated with QS that permits survival in the presence of toxic molecules, indicating paths for mutational complementation studies and inhibition of QS molecules by herbicides. This system represents a model of metabolic and physiological plasticity necessary for adaptation to agricultural environments. This model has the potential to be applied to biofilms produced by several species, thereby increasing the potential for degradation of xenobiotics and resistance to environmental stresses.

## Acknowledgement

The authors thank Marco Aurelio Sigismondi Ahuaji Filho for assistance with the resolution of the figures.

## S1 Appendix Caption

S1 - Analytical parameters unique to each analyte to configure MRM for each individual analyte.

